# Neural response properties predict perceived contents and locations elicited by intracranial electrical stimulation of human auditory cortex

**DOI:** 10.1101/2023.05.06.539680

**Authors:** Qian Wang, Lu Luo, Na Xu, Jing Wang, Ruolin Yang, Guanpeng Chen, Jie Ren, Guoming Luan, Fang Fang

## Abstract

Intracranial electrical stimulation (iES) of auditory cortex can elicit sound experiences with a variety of perceived contents (hallucination or illusion) and locations (contralateral or bilateral side), independent of actual acoustic inputs. However, the neural mechanisms underlying this elicitation heterogeneity remain undiscovered. Here, we collected subjective reports following iES at 3062 intracranial sites in 28 patients and identified 113 auditory cortical sites with iES-elicited sound experiences. We then decomposed the sound-induced intracranial electroencephalogram (iEEG) signals recorded from all 113 sites into time-frequency features. We found that the iES-elicited perceived contents can be predicted by the early high-γ features extract from sound-induced iEEG. In contrast, the perceived locations elicited by stimulating hallucination sites and illusion sites are determined by the late high-γ and long-lasting α features, respectively. Our study unveils the crucial neural signatures of iES-elicited sound experiences in human and presents a new strategy to hearing restoration for individuals suffering from deafness.

## Introduction

We gaze upon the glimmers of light; we hark the whispers of sound. Yet, what if the resplendence and the harmonies dost fade into the abyss?

A general framework for perception formation is that the activity of neurons in sensory cortex translates the external world into internal experiences. This framework suggests that it is possible to generate an artificial percept by activating sensory cortical neurons without altering the physical reality. More than half a century ago, using intracranial electrical stimulation (iES) in neurosurgical patients, Penfield and colleagues revealed that artificial percepts can be generated by activating sensory cortex without altering the external world^1, 2, 3^. When superior temporal cortex was stimulated, patients can experience a remarkable variety of perceived contents (hallucination: perceptions of sounds that do not exist, e.g. “buzzing”, “water dropping” or “familiar music”; illusion: altered perceptions of existing sounds, e.g. muffling of the environment sounds) and locations (contralaterally, ipsilateral, or bilaterally to the iES side)^2, 4, 5, 6, 7^. These findings introduce a basic framework of auditory cortical prosthesis techniques for the reconstruction of hearing capabilities in individuals suffering from peripheral deafness^8, 9, 10, 11^. However, the limited understanding of neural mechanisms underlying the diverse iES-elicited auditory percepts currently constrains the progress of these techniques.

Neurons in the auditory cortex do not respond to acoustic stimuli uniformly. Instead, their response properties exhibit variability in feature selectivity, temporal profile, and excitability^12, 13, 14, 15, 16, 17^. An appealing assumption posits that the diversity of neuronal response properties corresponds to the heterogeneity of iES-elicited auditory percepts. This assumption is supported by findings indicating that auditory cortical sites with iES-elicited hallucinations and those with iES-elicited illusions exhibit distinct anatomical distributions^7, 18, 19, 20^. However, other studies have reported overlapping distributions of hallucination and illusion sites within the auditory cortex^5, 6^. These inconsistent findings make it challenging to create a uniform anatomical distribution map of subjective sound experiences, rendering iES-elicited percepts unpredictable. Therefore, a more direct method to examine this assumption involves determining if any intrinsic response signatures can predict the perceived contents and locations of iES-elicited percepts; however, there is currently a lack of empirical evidence in this regard.

To investigate whether neuronal populations that engage in distinct iES-elicited percepts encode and represent sound information differently, we combined iES and intracranial electroencephalogram (iEEG) approaches in the human auditory cortex with high spatiotemporal precision. In the current study, we classified the cortical sites based on the contents and locations of iES-elicited sound experiences and then extracted the critical neuronal properties from iEEG signals which can predict the type of iES-elicited percepts. The iES approach enabled us to reveal perceived contents and locations of the internal sound experience when a neuronal population around an electrode site was stimulated. Meanwhile, the iEEG approach elucidated how the same neuronal population encodes and represents external acoustic stimuli. The combination of these two approaches provided a rare and unique opportunity to explore the relationship between intrinsic neuronal properties and perceived content and location^21, 22, 23^.

## Results

We collected first-person reports elicited by iES applied to 3062 intracranial sites in 28 epileptic patients who underwent stereo-electrode implantation to identify the source of their seizure onset zones. The iES procedure was conducted by clinicians who were blinded to the study’s objectives, and the first-person reports were recorded. The reports were examined and screened by an experienced neurologist (Dr. Jing Wang). Our analysis revealed that subjective reported effects were found in 1316 sites (42.98%). Among these sites, 153 exhibited iES-elicited auditory percepts, which were referred to as auditory elicitation sites. We then identified 113 auditory elicitation sites located in the auditory cortex, including Heshl’s gyrus (HG, n = 79), middle superior temporal gyrus (mSTG, n = 16), and posterior superior temporal gyrus (pSTG, n = 18). According to previous studies, HG encompasses the major part of the primary auditory cortex, while mSTG and pSTG are considered mostly as non-primary auditory cortex^24, 25^.

We classified auditory elicitation sites based on the perceived contents and locations elicited by iES. We identified two subsets of auditory elicitation sites: hallucination (n = 77) and illusion (n = 36) sites (**Figure 1A**), with hallucinations defined as generative perceptions of sounds that do not exist acoustically, and illusions defined as altered perceptions of sounds that do exist acoustically. Most of the hallucination sites exhibited simple sound contents (n = 74), while three sites exhibited complex sound contents such as familiar music and songs ^6, 7^. Illusion sites were further divided into suppression (n = 12) and echo (n = 24) sites^5, 6^. As shown in **Figure 1A**, we classified auditory elicitation sites based on their perceived locations, with most hallucination (n = 53) and illusion (n = 22) sites exhibiting perceived locations contralateral to their iES sides. A small subset of hallucination (n = 13) and illusion (n = 8) sites exhibited perceived locations bilaterally (e.g. “‘Cheep’ sounds in both ears appear.”). Patients failed to report the exact perceived locations of the iES-elicited auditory percepts in 17 sites (hallucination, n = 11; illusion, n = 6). We listed detailed subjective reports in **Table S2**. All the perceived contents and locations that we observed in the current study have been reported in previous studies^5, 6, 7, 26, 27^.

**Figure 1.**
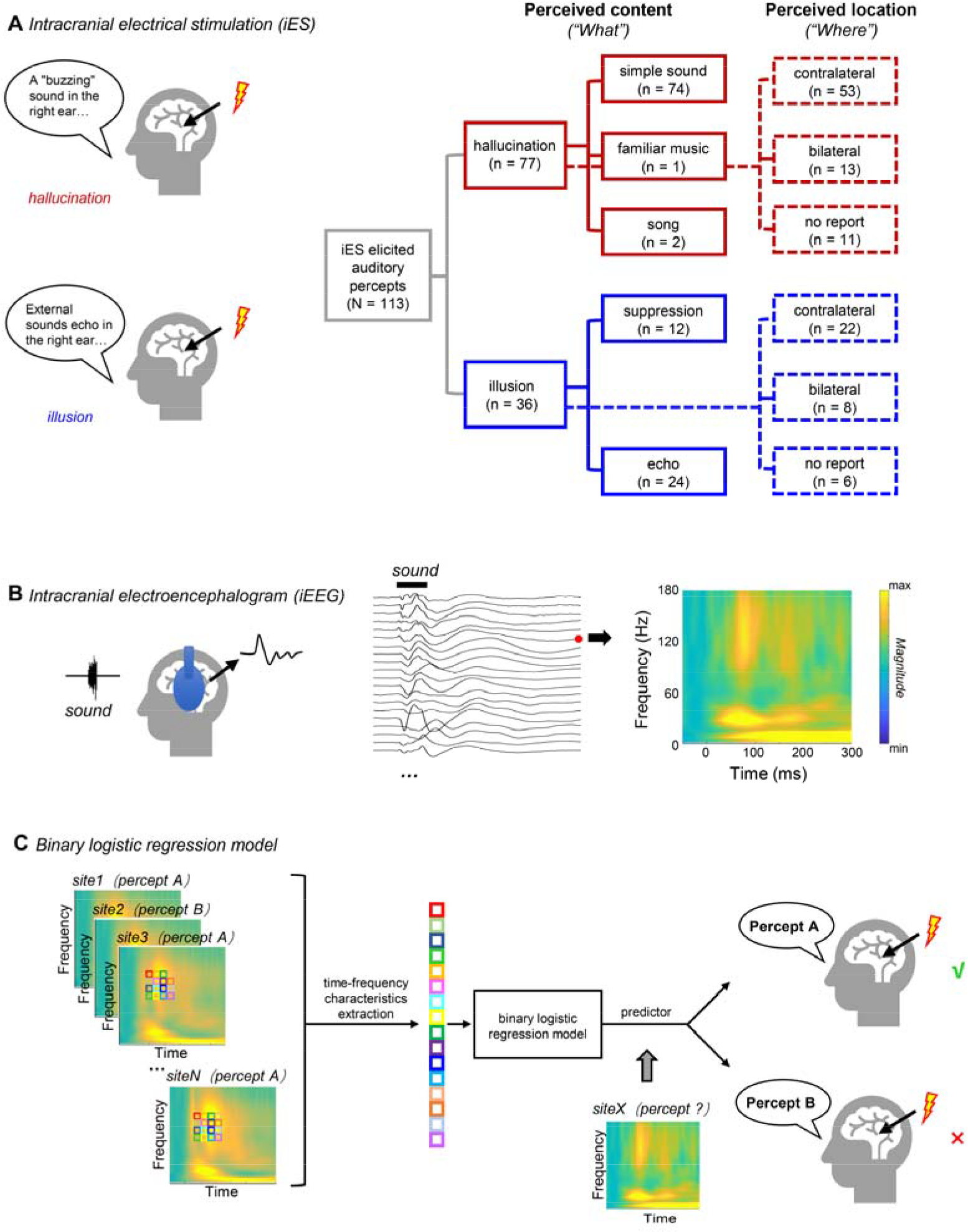
Experimental procedures and regression model. **(A)** Schematic illustration of the iES procedure (*left*) and the tree diagram of the classification of iES-elicited auditory percepts (*right*). *Black dialogue boxes* present two representative self-reports of iES-elicited auditory percepts. *Red boxes* indicate hallucinations. *Blue boxes* indicate illusions. *Solid boxes* represent the classification based on perceived contents. *Dashed boxes* represent the classification based on perceived locations. **(B)** Schematic illustration of the iEEG procedure (*left*), sound-induced responses (*middle*), and the time-frequency map of a representative sound-induced response (*right*). **(C)** Schematic illustration of the binary logistic regression model.

In addition to the iES procedure, patients also underwent a passive listening task during which their iEEG signals were continuously recorded (**Figure 1B**). Binaural white noises were presented to all 28 patients and the sound-induced iEEG signals were analyzed to investigate the intrinsic response properties of the neuronal population surrounding each site. In order to explore how auditory space was represented by the neuronal population around each site, 22 patients (P236-P421) were presented with ipsilateral and contralateral sounds (details are provided in the **Methods** section).

### Sound induced high-**γ** band signatures predicted perceived contents elicited by iES

We explored the anatomical distribution of hallucination and illusion sites. Figure 2B shows that more hallucination (72.70%) and illusion (63.9%) sites were localized in HG, with relatively smaller proportions localized in mSTG and pSTG. Regarding the proportions of hallucination sites and illusion sites, there was no significant difference among HG, mSTG, and pSTG (χ2 (2) = 1.590; *p* = 0.452). However, we observed a hemispheric asymmetry, with more illusion sites localized in the left hemisphere (69.4%) (χ2 (1) = 6.941; *p* = 0.008) ^6^.

**Figure 2.**
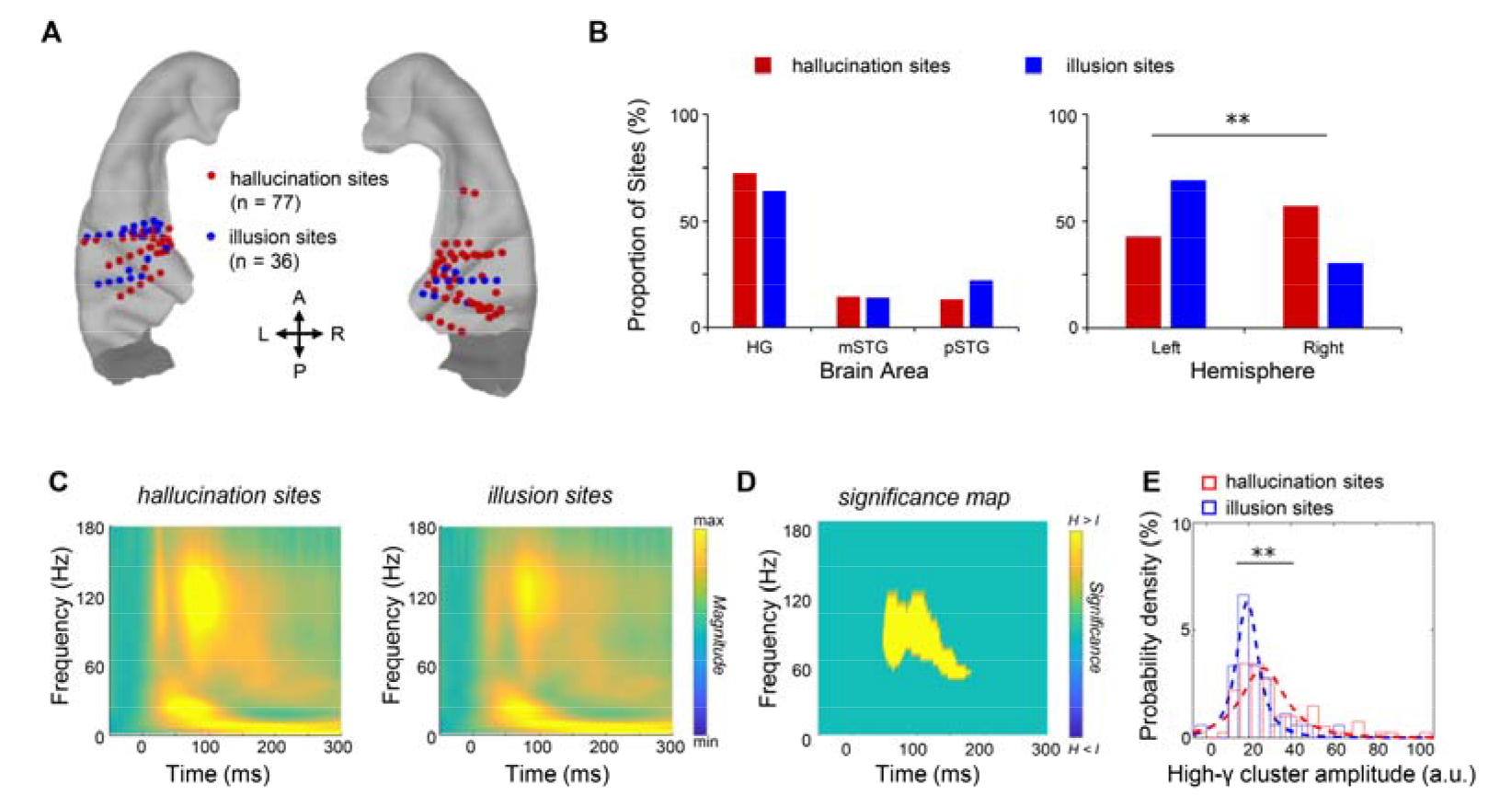
Anatomical and time-frequency characteristics of hallucination and illusion sites. **(A)** Hallucination (*red*) and illusion (*blue*) sites are shown on the temporal lobe of the ICBM152 brain template. **(B)** Proportions of hallucination (*red*) and illusion (*blue*) sites in HG, mSTG, and pSTG (*left*) and in the left and right hemispheres (*right*). **(C)** Averaged time-frequency maps of sound-induced responses of hallucination sites (*left*) and those of illusion sites (*right*). **(D)** The statistical parametric map of time-frequency differences between sound-induced responses of hallucination sites and those of illusion sites. *Yellow areas* indicate significant time-frequency clusters (*p* < 0.005, *FDR* corrected). **(E)** Comparison of high-γ cluster (50 to 116 Hz; 49 to 139 ms) amplitudes between hallucination and illusion sites. *H: hallucination; I: illusion.* ** *p* < 0.01.

We proceeded to compare the time-frequency map of the sound-induced responses in hallucination and illusion sites (Figure 2C). Our results revealed that the hallucination sites showed an early increase in power in the high-γ band (50 to 116 Hz) in response to the sounds, relative to the illusion sites (*p* < 0.005, *FDR* corrected) (Figure 2D). Additionally, we observed that the mean amplitude of the early high-γ cluster was significantly larger in the hallucination sites than that in the illusion sites (*t* (111) = 3.005, *p* = 0.003; Figure 2E). One possible explanation for this difference in high-γ amplitude is that the neuronal excitability of the hallucination sites is higher than that of the illusion sites. Here, we estimated the excitability of each site by defining the elicitation threshold as the minimum current intensity required to elicit a conscious percept ^28^. We found no significant difference between the averaged elicitation threshold of the hallucination sites (2.12 ± 1.22 mA; mean ± SD) and that of the illusion sites (2.07 ± 0.85 mA) (*t* (111) = 0.267, *p* = 0.790), demonstrating that different iES-elicited perceived contents cannot be explained by the neuronal excitability difference. Taken together, these findings show that the neuronal population corresponding to hallucination is characterized by an enhancement of sound-induced early high-γ activities.

We then employed binary logistic regression to examine whether the high-γ signatures we observed could predict the perceived contents (hallucination or illusion) elicited by iES. As illustrated in Figure 1C, we constructed a binary regression model using the amplitudes of discrete time-frequency bins within the high-γ range (50 to 116 Hz, 49 to 139 ms) extracted from sound-induced iEEG signals (Figure 2D). Our analysis revealed that the high-γ band signatures extracted from iEEG responses to external sounds could predict 85.0% of the perceived contents (94.8% for hallucination, 63.9% for illusion) elicited by iES (χ2 (24) = 50.346, *p* = 0.001).

We further categorized the hallucination and illusion sites based on the details of their perceived contents elicited by iES (Figure 1A)^7^. We found that the majority of the hallucination sites (96.10%) exhibited simple sound perception. In contrast, a dichotomy was observed among illusion sites, with 33.33% of the sites showing suppression perception and 66.67% of the sites showing echo perception. Most suppression and echo sites were localized in HG, with relatively smaller proportions in mSTG and pSTG (Figure 3A and B). Regarding the proportions of the suppression and echo sites, there was no significant difference among HG, mSTG, and pSTG, suggesting a consistent distribution across the three areas (χ^2^ (2) = 2.010, *p* = 0.366). No hemispheric asymmetry was observed either (χ^2^ (1) = 1.047, *p* = 0.306).

**Figure 3.**
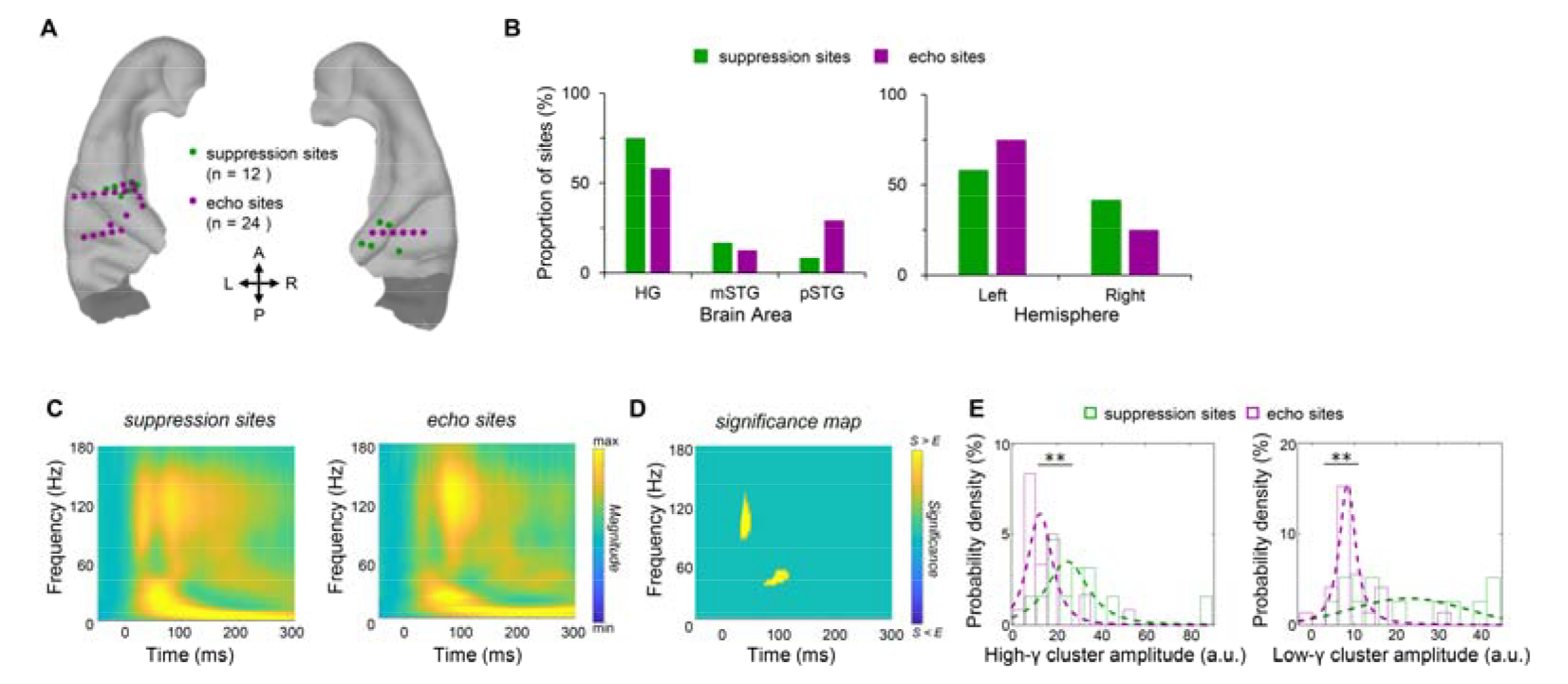
Anatomical and time-frequency characteristics of suppression and echo sites. **(A)** Suppression (*green*) and echo (*purple*) sites are shown on the temporal lobe of the ICBM152 brain template. **(B)** Proportions of suppression (*green*) and echo (*purple*) sites in HG, mSTG, and pSTG (*left*) and the left and right hemispheres (*right*). **(C)** Averaged time-frequency maps of sound-induced responses of suppression (*left*) and those of echo (*right*) sites. **(D)** The statistical parametric map of time-frequency differences between sound-induced responses of suppression sites and those of echo sites. *Yellow areas* indicate significant time-frequency clusters (*p* < 0.001). **(E)** Comparison of a high-γ cluster (*left*; 87 to 124 Hz; 31 to 47 ms) and a low-γ cluster (*right*; 38 to 50 Hz; 72 to 115 ms) amplitudes between suppression (*green*) and echo (*purple*) sites. *Error bars* represent the standard error across sites. ** *p* < 0.01. S, suppression; E, echo.

We then compared the time-frequency maps of sound-induced responses in suppression and those in echo sites (Figure 3C) and found that suppression sites showed an early high-γ activity (31 to 47 ms; 87 to 124 Hz) and a late γ activity (72 to 115 ms; 38 to 50 Hz) enhancement in response to sounds, relative to echo sites (*p* < 0.005, *FDR* corrected) (Figure 3D). Both amplitudes of time-frequency clusters were significantly larger for suppression sites than for echo sites (high-γ cluster: *t* (34) = 3.072, *p* = 0.004; γ cluster: *t* (34) = 3.146, *p* = 0.003; Figure 3E). We found no significant difference between the elicitation thresholds of suppression sites (1.79 ± 0.81 mA) and those of echo sites (2.21 ± 0.86 mA) (*t* (34) = 1.397, *p* = 0.171).

We built two binary regression models using the amplitudes of discrete time-frequency bins extracted from the high-γ range (87 to 124 Hz; 31 to 47 ms) and the γ range (38 to 50 Hz; 72 to 115 ms) (Figure 3D) to predict perceived contents (suppression or echo), respectively. We found that the high-γ band signatures could predict 83.3% of perceived contents (suppression: 66.7%; echo: 91.7%) (χ^2^ (9) = 19.102, *p* = 0.024), and the γ band signatures could predict 91.7% of perceived contents (suppression: 83.3%; echo: 95.8%) (χ^2^ (8) = 26.992, *p* = 0.001). Together, these results demonstrate that high-γ and γ responses extracted from sound-induced iEEG signals can serve as powerful predictors of the perceived contents elicited by iES.

### Intrinsic high-**γ** temporal profiles characterized iES-elicited perceived contents

Previous iEEG studies have shown that sound-induced high-γ responses can be classified into transient and sustained types based on their distinct temporal profiles^29, 30, 31, 32^, suggesting the existence of functional dichotomy within neuronal populations characterized by distinct temporal profiles^33^. We also found that the iES-elicited perceived contents, including simple sound hallucinations and suppression/echo illusions, were associated with different sound-induced high-γ responses. Therefore, we investigated whether the diversity of iES-elicited perceived contents could be explained by the distinction of high-γ temporal profiles.

We applied convex non-negative matrix factorization (NMF) to the sound-induced high-γ responses^34^. As shown in Figure 4A, we discovered two classes of responses that could explain 87.01% of the variance in the data: one with a strong response at the sound onset (transient), and the other with a long-lasting response (sustained). To compare the temporal properties of hallucination, echo, and suppression sites further, we calculated the relative weight of each site (details in Methods). For a given site, a higher relative weight value indicates a more transient response. Based on the comparison between the two NMF weights, we classified 27 transient sites and 86 sustained sites (**Figure 4B**). Figure 4C shows that 70.4% of transient sites and 67.4% of sustained sites were engaged in hallucination. Moreover, transient sites were more frequently involved in the perception of suppression (22.0%), compared to their involvement in echo perception (7.4%) (χ^2^ (2) = 7.719, *p* = 0.021). Figure 4D reveals that the relative weight of suppression sites was greater than that of echo sites (*t*(34) = 2.588, *p* = 0.014) and that of hallucination sites (*t*(87) = 1.93, *p* = 0.057). Additionally, we observed that the latency of suppression sites (21.16 ± 3.98 ms) was shorter than that of echo sites (41.59 ± 4.52 ms) (*t*(34) = 2.915, *p* = 0.006) and that of hallucination sites (32.92 ± 2.39 ms) (*t*(87) = 1.879, *p* = 0.064, *marginally significant*). These results indicate that the neuronal population engaged in suppression perception is characterized by rapid and transient sound-induced high-γ responses. Additionally, our findings elucidate the relationship between high-γ temporal profiles and subjective auditory experiences, offering empirical support for the functional dichotomy within neuronal populations possessing distinct temporal profiles, a concept hinted at by previous researches^29, 30, 31, 32^.

**Figure 4.**
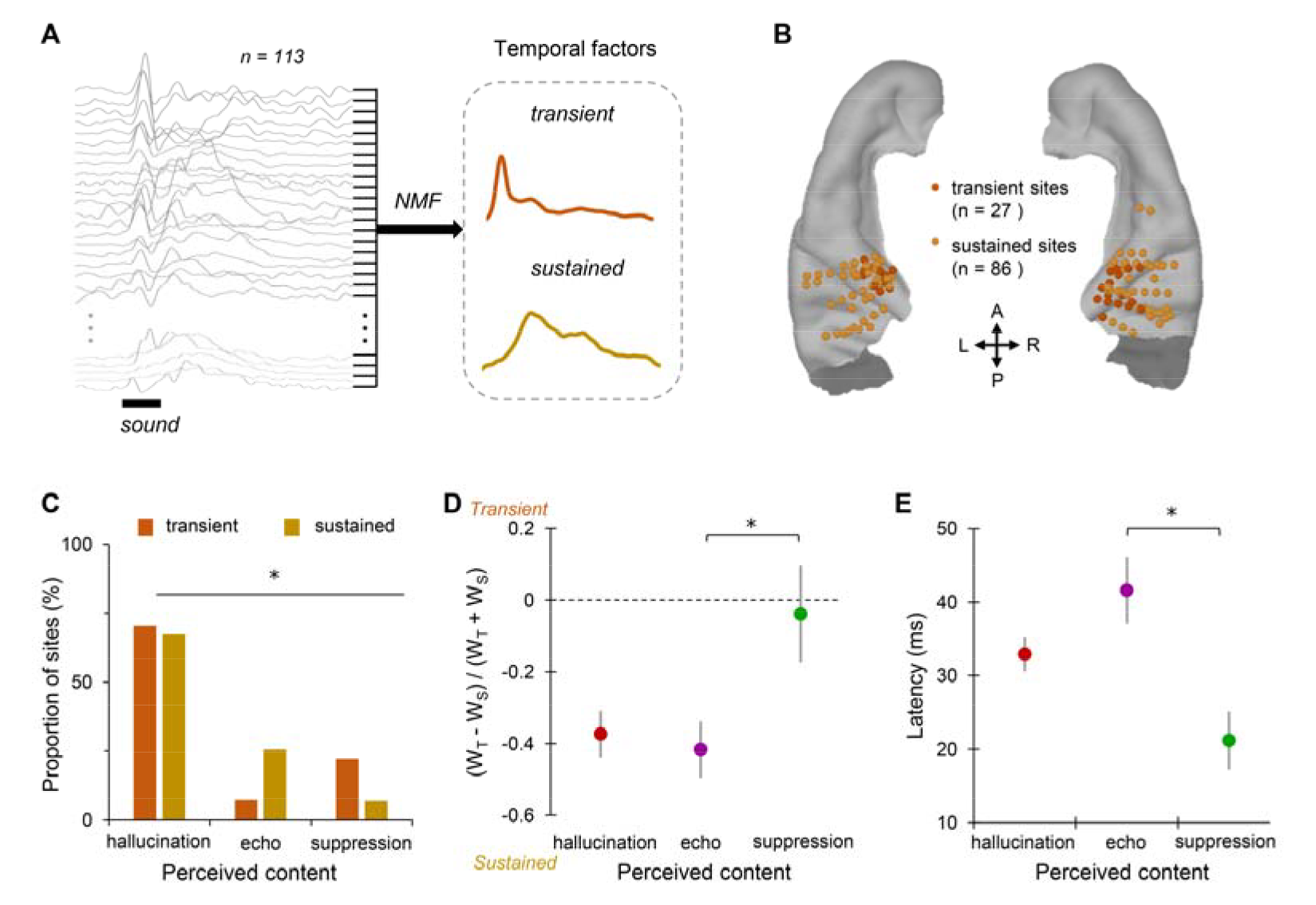
High-γ temporal profiles and latencies of hallucination, echo, and suppression sites. **(A)** NMF clustering results for *k* = 2. Averaged high-γ responses were normalized and then entered into the clustering pool (n = 113). Two temporal factors were observed. One factor showed a strong response at sound onset (Transient, *orange curve*), and the other showed a slow and sustained response (Sustained, *yellow curve*). **(B)** Spatial distributions of transient (*orange*) and sustained sites (*orange*). **(C)** Proportions of transient (*orange*) and sustained (*yellow*) sites exhibited iES-elicited hallucination, echo, and suppression. **(D)** Comparison of relative activation weights for sites exhibiting iES-elicited hallucination, echo, and suppression perception. **(E)** Averaged latencies of hallucination, echo, and suppression sites. *Error bars* represent the standard errors across sites. *W_T_*, transient weight; *W_S_,* sustained weight. * *p* < 0.05.

### Sound induced high-**γ** band signatures predicted perceived locations of hallucination sites

An iES-elicited auditory percept could be perceived either contralaterally or bilaterally to the stimulation side^6, 7^. We observed that most of the hallucination and illusion sites were perceived contralaterally, while a small proportion of hallucination and illusion sites were perceived bilaterally (Figure 1A).

To further investigate the differences between contralateral and bilateral hallucination sites, we analyzed their anatomical and time-frequency characteristics. As shown in Figures 5A and B, compared with bilateral hallucination sites, more contralateral hallucination sites were located in the HG (χ^2^ (2) = 11.349; *p* = 0.003). No hemispheric asymmetry was observed (χ^2^ (1) = 0.307; *p* = 0.579). Regarding time-frequency characteristics, we found that contralateral hallucination sites showed a late amplitude enhancement (107 to 158 ms) in the high-γ band (87 to 153 Hz) in response to sounds compared to bilateral hallucination sites (*p* < 0.005, *FDR* corrected) (Figures 5C and D). The mean amplitude of the high-γ cluster of contralateral hallucination sites was significantly larger than that of bilateral hallucination sites (*t* (64) = 2.732, *p* = 0.008; Figure 5E). No difference was found between the elicitation thresholds of contralateral hallucination sites (2.06 ± 1.21 mA) and bilateral hallucination sites (2.31 ± 1.30 mA) (*t* (64) = 0.662, *p* = 0.510).

**Figure 5.**
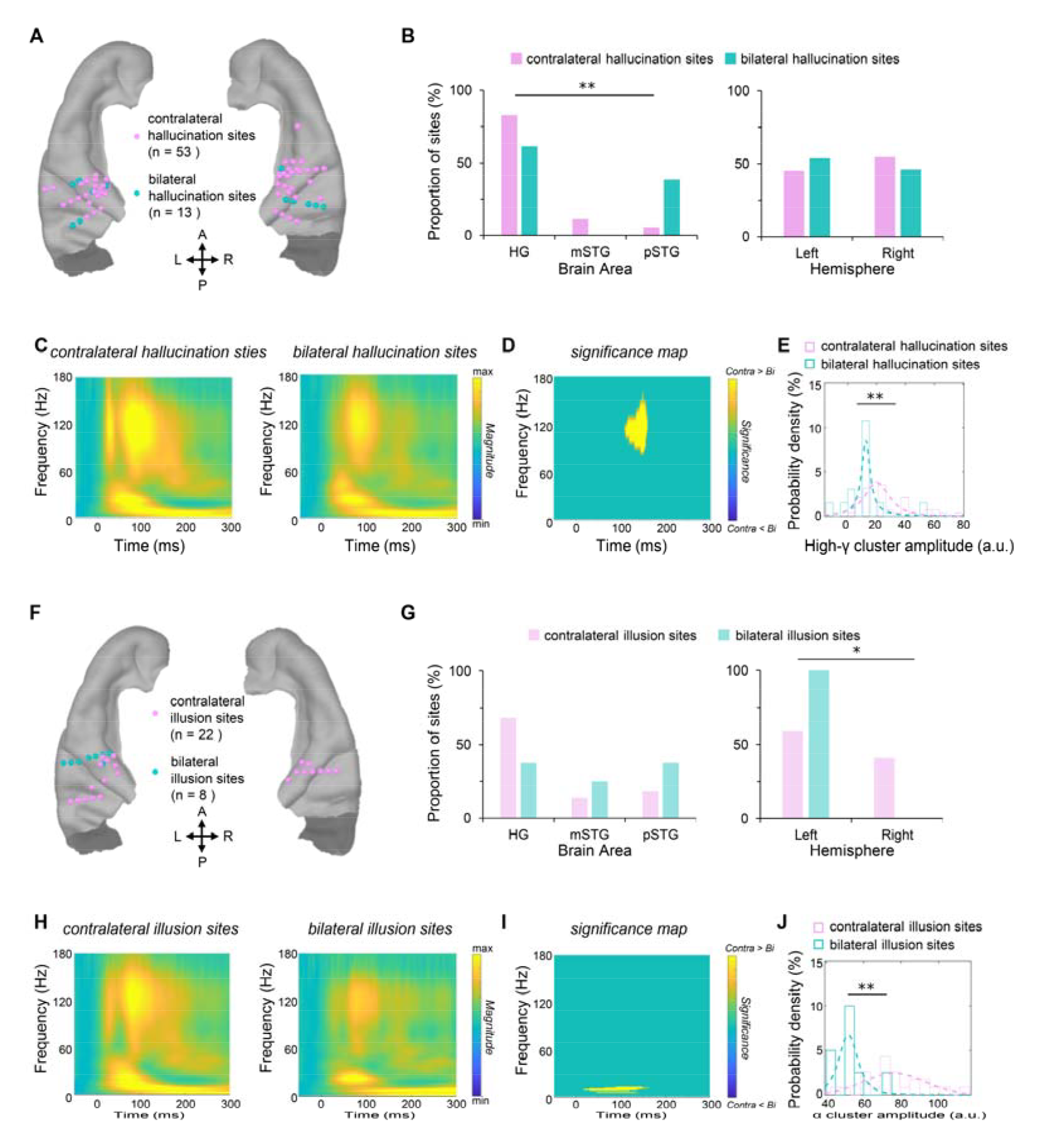
Anatomical and time-frequency characteristics of sites with different iES-elicited perceived locations. **(A)** Contralateral (*pink*) and bilateral (*cyan*) hallucination sites are shown on the temporal lobe of the ICBM152 brain template. **(B)** Proportions of contralateral (*pink*) and bilateral (*cyan*) hallucination sites in HG, mSTG, and pSTG (*left*) and the left and right hemispheres (*right*). **(C)** Averaged time-frequency maps of the sound-induced responses of contralateral hallucination sites (*left*) and those of bilateral hallucination sites (*right*). **(D)** The statistical parametric map of time-frequency differences between the sound-induced responses of contralateral hallucination sites (*left*) and those of bilateral hallucination sites (*right*). **(E)** Comparison of the high-γ cluster (87 to 153 Hz; 107 to 158 ms) amplitudes between contralateral (*pink*) and bilateral (*cyan*) hallucination sites. **(F)** Contralateral (*pink*) and bilateral (*cyan*) illusion sites are shown on the temporal lobe of the ICBM152 brain template. **(G)** Proportions of contralateral (*light pink*) and bilateral (*light cyan*) illusion sites in HG, mSTG, and pSTG (*left*) and the left and right hemispheres (*right*). **(H)** Averaged time-frequency maps the sound-induced responses of contralateral illusion sites (*light pink*) and those of bilateral illusion sites (*light cyan*). **(II)** The statistical parametric map of time-frequency differences between the sound-induced responses of contralateral illusion sites (*left*) and those of bilateral illusion sites (*right*). **(J)** Comparison of the α cluster (7 to 15 Hz; 17 to 164 ms) amplitudes between contralateral (*light pink*) and bilateral (*light cyan*) illusion sites. *Error bars* represent the standard error across sites. *Yellow areas* indicate significant time-frequency clusters (*p* < 0.005, *FDR* corrected). *Contra: contralateral; Bi: bilateral.* * *p* < 0.05; ** *p* < 0.01.

These results suggest that the neuronal population corresponding to contralateral hallucination is characterized by a late high-γ power enhancement in response to acoustic sounds, mainly located in the primary auditory cortex.

We constructed a binary regression model using the amplitudes of discrete time-frequency bins extracted from the high-γ range (87 to 153 Hz; 107 to 158 ms) shown in Figure 5D. Our analysis revealed that the perceived location could be predicted with an accuracy of 83.3% (χ^2^ (9) = 18.490, *p* = 0.030).

### Sound induced **α** band signatures predicted perceived locations of illusion sites

We conducted analysis to investigate the anatomical and time-frequency characteristics of contralateral and bilateral illusion sites. Regarding the proportions of the contralateral illusion sites and the bilateral illusion sites, As shown in Figures 5F and G, there was no significant difference among HG, mSTG, or pSTG (χ^2^ (2) = 2.313, *p* = 0.315). However, we found an asymmetry effect only in the bilateral illusion sites (left: 100%; right: 0%), but not in the contralateral illusion sites (left: 59.1%; right: 40.9%) (χ^2^ (1) = 4.675; *p* = 0.031).

We also examined the time-frequency characteristics of contralateral and bilateral illusion sites. The contralateral illusion sites exhibited an early amplitude increase (17 to 164 ms) in the α band (7 to 15 Hz) in response to the sounds, relative to the bilateral illusion sites (*p* < 0.005, *FDR* corrected) (Figures 5H and I). The amplitude of the α cluster was significantly larger for the contralateral illusion sites than those for bilateral illusion sites (*t*(28) = 3.626, *p* = 0.001; Figure 5J). No significant difference was found between the elicitation threshold of contralateral illusion sites (2.27 ± 0.92 mA) and that of bilateral illusion sites (2.06 ± 0.68 mA) (*t*(28) = 0.587, *p* = 0.562). These results suggest that the neuronal population corresponding to contralateral illusion is characterized by an alpha power enhancement in response to sounds.

A binary regression model using the amplitudes of discrete time-frequency bins extracted from the α range (7 to 15 Hz; 17 to 164 ms; Figure 5I) predicted perceived locations with an accuracy of 90.0%, with 90.9% accuracy for contralateral and 87.5% accuracy for bilateral perception (χ^2^ (3) = 20.793, *p* < 0.001).

Overall, our findings suggest that the high-γ and α band signatures extracted from sound-induced iEEG can be used to predict perceived locations of hallucination and illusion sites, respectively.

### Auditory space was differently represented in hallucination and illusion sites

Our findings that the perceived location elicited by stimulating hallucination sites and that of illusion sites could be determined by the sound-induced high-γ and α activities raise an interesting question: whether auditory space is represented differently in hallucination and illusion sites? To address this question, we presented ipsilateral and contralateral sounds to 22 patients (P236-P421) and analyzed a total of 77 sites divided into four groups (contralateral hallucination, bilateral hallucination, contralateral illusion, and bilateral illusion) (Figure 6A). Time-frequency analyses were performed to extract spectra-temporal features in the high-γ and α bands for each site (Figure 6B). We then trained and decoded the auditory space with single-trial spectra-temporal features using support vector machine approaches via leave-one-out cross-validations. First, we trained and decoded the auditory space with single-trial spectra-temporal features in the high-γ band. Second, we trained and decoded the auditory space with the α band features. To compare the decoding accuracies among site groups, 2 (Perceived Location: contralateral, bilateral) × 2 (Perceived Content: hallucination, illusion) ANOVAs were performed for both high-γ and α models.

**Figure 6.**
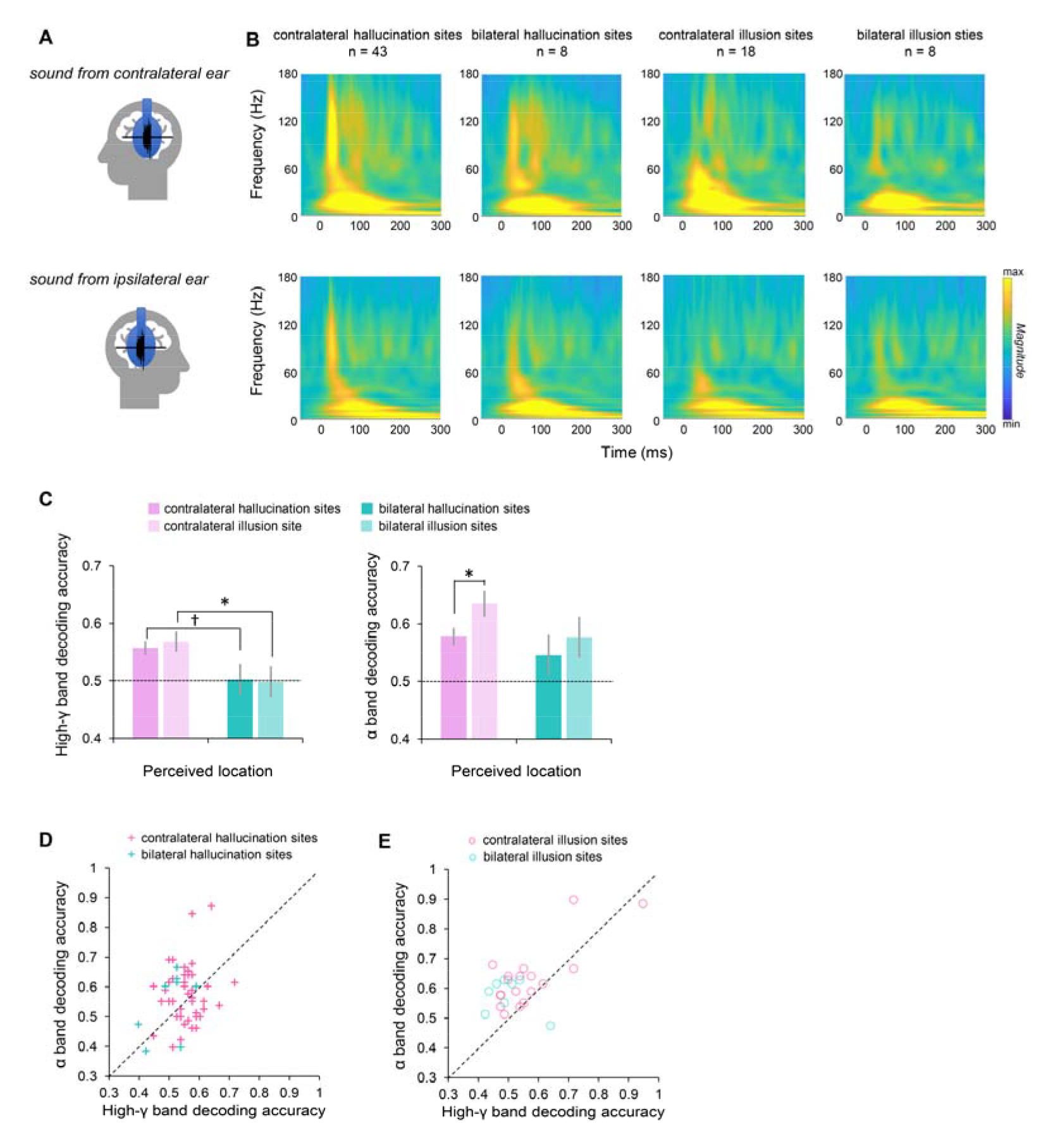
Decoding accuracy of sites with different iES-elicited percepts. **(A)** Sounds were present either contralaterally or ipsilaterally. **(B)** Averaged time-frequency maps of neural signals responded to contralateral (*up*) and ipsilateral (*down*) sounds. **(C), (D), (E)** Comparisons of decoding accuracies of the high-γ (*left*) and α (*right*) models. *, *p* < 0.05; †, *p* < 0.1.

For the decoding accuracy using the high-γ band features, we found a significant main effect of Perceived Location (*F*(1,73) = 7.921, *p* = 0.006), indicating that auditory space was represented differently between contralateral and ipsilateral sounds. No significant effect of Perceived Content or interaction effect was found. Post hoc tests confirmed the Perceived Location effect in illusion conditions (contralateral hallucination vs. bilateral hallucination: *p* = 0.067; contralateral illusion vs. bilateral illusion: *p* = 0.038, Figure 6C, left). For the α model, we found marginal significant main effects of Perceived Location (*F*(1,73) = 2.605, *p* = 0.111) and Perceived Content (*F*(1,73) = 2.436, *p* = 0.123). No significant interaction effect was found. Post hoc tests showed that the decoding accuracy of contralateral illusion sites was higher than contralateral hallucination sites (*p* = 0.041, Figure 6C, right).

We also compared the decoding accuracy using the high-γ and α band features in each site group. We found that the decoding accuracy using the α band features was higher than that of the high-γ band features in contralateral illusion (*t*(17) = 3.668, *p* = 0.002) and bilateral illusion (*t*(7) = 2.126, *p* = 0.071)) sites (Figure 6E). No significant difference was found in contralateral hallucination or bilateral hallucination sites (both *p* > 0.5; Figure 6D). These results suggest that the α band activity plays an essential role in representing auditory space in illusion sites.

## Discussions

The elicitation of subjective sound experiences through electrical stimulation of the auditory cortex has been known for over half a century, yet the underlying mechanisms remain poorly understood. In this study, we employed a combination of iES and iEEG techniques to investigate the relationship between iES-elicited sound experiences and neuronal response properties in the human auditory cortex. By categorizing iES-elicited sound experiences based on their perceived contents (hallucination and illusion) and locations (contralateral or bilateral side), we were able to further categorize illusion sites into echo and suppression subsets. Our results showed that sites with diverse perceived contents were determined by the amplitudes, temporal profiles, and latencies of sound-induced high-γ band activity. Notably, we observed that the perceived locations of hallucination sites and those of illusion sites could be predicted by the high-γ band and α band activities, respectively. Furthermore, we demonstrated that auditory space information could be decoded from high-γ and α band time-frequency features at the single-site level, with the α band features primarily contributing to the decoding performance of illusion sites.

The variability of iES effects has been previously linked to stimulation patterns^35^. To address this issue, we conducted a strict control of stimulation parameters, including frequency, pulse width, and stimulation duration, during each iES trial in the present investigation. Our sole alteration of current intensity allowed for the evaluation of the elicitation threshold of each site, and we systematically compared the thresholds across sites with different perceived contents and locations. Notably, our results showed no significant difference among the elicitation thresholds of these sites, which suggests that the observed diversity in iES-elicited sound experiences cannot be solely attributed to the variation in stimulation patterns.

Using the classical "what and where" framework^36, 37, 38^, we classified the iES-elicited sound experiences based on their perceived contents and locations. All the iES-elicited perceived content and locations has been previously reported in literature^1, 2, 3, 4, 5, 6, 7^.

On one hand, iES-elicited perceived contents can be classified into hallucination and illusion categories^6, 7, 18, 20^. Hallucination is a common iES-elicited effect that contains simple (elementary sounds) and complex (music or voice) subcategories^6, 7^, while illusion is relatively rare and contains suppression (hearing sounds less clearly or from a distance) and echo subcategories^5, 6, 39^. Using time-frequency analysis of intracranial signals^26, 40^, the current study established a relationship between high-γ band responses in the auditory cortex and perceived contents. Previous studies have revealed that high-γ responses can be distinguished by their temporal response profiles: transient and sustained profiles^29, 30, 32^. This neurophysiological dichotomy in the high-γ temporal profile was assumed to be functionally related to the distributed processing of auditory information^33^, but empirical evidence for this relationship was lacking. Our findings supported this assumption by revealing that sites with different temporal response properties correspond to different perceived contents. Specifically, transient responses would be consistent with the motor-induced suppression of the auditory cortex, which is involved in perceptual guidance of action^41, 42^, while sustained responses would be consistent with feedback influences on auditory object recognition processes^33^.

On the other hand, both hallucinations and illusions can be perceived contralaterally or bilaterally, based on their iES-elicited perceived locations^5, 6^. The current study established a relationship between high-γ and α band responses in the auditory cortex and perceived locations. Previous studies have found that both high-γ and α band signals can encode and characterize perceptual space^43, 44^. Interestingly, in the current study, we found that hallucination sites and illusion sites encoded auditory space via high-γ and α band signals, respectively. We further revealed that auditory space information could be decoded from high-γ and α band time-frequency features, withα band features predominantly contributing to the decoding performance of illusion sites. This finding is consistent with a previous research that has shown that the α band plays an important role in illusions^45, 46^.

Furthermore, our study provides novel insights into the neural mechanisms underlying the formation of auditory illusions. The debate regarding the contributions of feedforward and feedback signals in illusion formation has been a major point of contention in the field^16, 47, 48^. According to the hierarchical model, primary sensory cortex is responsible for hallucinations while high-order cortex is responsible for illusions^5, 20, 49^. However, our results demonstrate that the distribution of illusion and hallucination sites along the auditory cortical hierarchy does not support this perspective. Instead, we found that a subset of illusion sites, specifically those responsible for suppression, exhibited a approximated response latency as that of the human primary auditory cortex^50^. Therefore, our findings support the idea that feedforward signals may play a primary role in generating illusions by directly modulating subjective experiences in humans.

In sum, our study revealed that the neuronal response properties of different sites in the auditory cortex can effectively predict subjective sound experiences elicited by iES. This finding holds promising approach for the development of auditory cortical prostheses for individuals with acquired deafness. Additionally, our results may provide a novel direction for the treatment of other auditory-related neurological and psychiatric disorders, such as tinnitus and schizophrenic hallucinations.

## Methods

### Participants

A cohort of 28 patients (10 females, age range 13-43 years) with drug-resistant epilepsy who underwent invasive stereo-electroencephalogram monitoring for potential surgical interventions at the Sanbo Brain Hospital of Capital Medical University (Beijing, China) were recruited. Patient selection criteria required the presence of at least one electrode implanted in the superior temporal gyrus. All participants were right-handed and self-reported normal hearing, and provided written informed consent for their participation. The study procedures were approved by the Ethics Committee of the Sanbo Brain Hospital of Capital Medical University and the Human Subject Review Committee of Peking University. Details of patient demographics are provided in Table S1.

### Stereo-electrode site localization and selection

All 28 patients were implanted with stereo-electrodes. Each stereo-electrode had 8-16 sites (0.8 mm in diameter, 2 mm in length, spacing 3.5 mm apart; Huake Hengsheng Medical Technology Co. Ltd., Beijing, China). The electrode implantations were determined based on clinical reasons. For stereo-electrode site localization, the post-implantation CT images were co-registered with the pre-implantation T1-weighted MRI scans for each subject using the SPM12 toolbox (available at https://www.fil.ion.ucl.ac.uk/spm/software/spm12/)^51^. Then, we identified individual electrodes on the aligned CT images and calculated the coordinates of electrode sites using the Brainstorm toolbox (available at http://neuroimage.usc.edu/brainstorm)^52^. Three-dimensional brain surfaces were reconstructed by pre-implantation MR images (T1-weighted or contrast-enhanced) and registered to the USC Brain atlas^53^ using the BrainSuite software (http://brainsuite.org/). The localization of each site was identified using the individual atlas of each patient. The sites localized in the Heshl’s gyrus (HG), middle superior temporal gyrus (mSTG), and posterior superior temporal gyrus (pSTG), were selected as auditory cortical sites. For visualization, we computed MNI coordinates of each site individually and displayed them on a cortical template (ICBM152). The sites with over-threshold impedances (> 50 kΩ), artifacts, and/or within epilepsy foci were excluded from data analyses.

### Intracranial electrical stimulation (iES)

All patients underwent iES mapping as part of a routine clinical procedure to determine essential sensory, motor, and other cognitive function using Nicolet® Cortical Stimulator (Natus Neuro, USA). In each iES trial, a train of biphasic electrical pulses (square-wave, 0.5 to 6 mA, 50 Hz, 200-μs pulse width, 5-s duration) was delivered to pairs of adjacent sites. During the iES procedure, patients were asked to perform a number counting task (speak loudly from one to one hundred) and to report immediately if they had any feelings. Patients were unaware of the timing of the stimulation and the anatomical location of the stimulation sites. Patients’ self-reports were reviewed and further classified by an experienced neuropsychologist (Dr. Jing Wang).

### Intracranial electroencephalogram (iEEG)

Intracranial EEG signals were recorded at a sampling rate of 512 Hz using a Nicolet video-EEG monitoring system (Thermo Nicolet Corp., USA) without any online filtering. Both the reference and ground electrodes were placed at the forehead of the patients. All further processing was performed offline.

To obtain the sound-induced responses, we recorded intracranial signals via stereo-electrode sites when patients were passively listening to Gaussian wideband noise bursts. The wideband-noise stimulus was synthesized in MATLAB environment (MathWorks, Natick, MA, USA) at the sampling rate of 48 kHz with 16-bit amplitude quantization and low-pass filtered at 10 kHz. The duration of the noise burst stimulus was 50 ms including the 5-ms linear ramp and damp. The interstimulus interval (ISI) randomly varied between 900 and 1100 ms. The acoustic stimulus was transferred using Creative Sound Blaster X-Fi Surround 5.1 Pro (Creative Technology Ltd, Singapore) and presented to patients with insert earphones (ER-3, Etymotic Research, Elk Grove Village, IL) at the sound pressure level of 65 dB SPL. Calibration of the sound level was carried out with the Larson Davis Audiometer Calibration and Electroacoustic Testing System (AUDit and System 824, Larson Davis, Depew, NY).

All patients were presented with binaural noise bursts (180 trials), and 22 patients (P236-P421) were additionally presented with ipsilateral (160 trials) and contralateral (160 trials) noise bursts. Patients were reclining in a ward in the hospital during the experiment. The experimental procedure was suspended for at least 2 hours after a seizure (if there was any), to avoid the seizure-induced cortical suppression effect.

### Preprocessing of intracranial signals

All the intracranial signals were pre-processed using the EEGLAB toolbox (Delorme and Makeig, 2004) in the MATLAB environment. The continuous signals were then segmented into epochs from −100 to 600 ms around the sound onset and normalized to the baseline (−100 to 0 ms). The epochs that contained interictal discharges or artifacts were rejected.

### Time-frequency analyses

Time-frequency analyses were performed on the epoched signal using continuous Morlet wavelet transformation (function *cwt*) in Matlab. For each electrode site, time-frequency maps with a frequency range from 2 to 180 Hz and a time range from −50 to 300 ms were calculated for each trial. The amplitudes of the time-frequency maps were then averaged across trials. Paired *t*-tests were performed point-by-point to compare the time-frequency components between two conditions. Significant time-frequency points (*p* < 0.005, two-tailed) were then clustered based on spatial adjacency, with a minimum of ten adjacent points to form a cluster. The temporal profiles in the high-γ band (60 to 140 Hz) and the alpha band (6 to 14 Hz) activities were subsequently extracted from the same time–frequency decomposing analysis.

### Unsupervised clustering of high-***γ*** temporal profiles

Previous studies have revealed that the temporal profiles of sound-induced high-γ waveforms can be divided into transient and sustained types, which may be engaged in different functions of auditory processing^29, 30, 32^. To examine whether these two types of temporal profiles differed and engaged in different iES-elicited percepts, we used convex non-negative matrix factorization (NMF) ^34^ to cluster the time series of sound-induced high-γ responses (−50 to 300 ms). The time series X [time points × sites] were estimated using the following function:

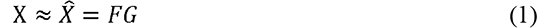

For a given site, the *G* matrix [clusters × sites] represents the clustering weight and the *F* matrix [time points × clusters] represents the prototypical time series of a given cluster.

We performed the clustering analysis for the averaged high-γ responses of 113 sites across subjects, which resulted in the time series X [180 time points × 113 sites]. To evaluate the explanatory power of the cluster results, we calculated the percent variance (*R*^2^) with the following function:

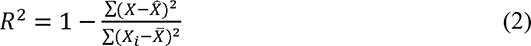

We calculated the silhouette index (SI) to evaluate the clustering goodness of fit using the *silhouette* function in Matlab. To test the statistical significance of the SI, the classification types for all sites were shuffled 1000 times to create the chance levels of the SI.

For NMF clustering results, to quantify the relative contribution of the transient and sustained components, we calculated a relative weight as:

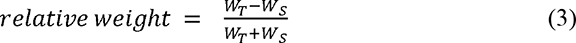

The resulting relative weight ranges from −1 to 1. A higher relative weight value indicates a more transient property.

### Spectra-temporal decoding

Based on the iES-elicited percepts, we divided the electrode sites into four groups: contralateral hallucination (n = 43), bilateral hallucination (n = 8), contralateral hallucination (n = 18), and bilateral hallucination (n = 8) (Figure 6). For each site, we built two models using the spectra-temporal features in the high-γ band (60 to 140 Hz) or those in the α band (6 to 14 Hz) respectively, to classify the neural activities induced by ipsilateral and contralateral sounds. We employed a support vector machine (SVM) approach for each model using single-trial spectra-temporal features ^30^.

The spectra-temporal features were extracted from 52 bins [13 spectra-steps × 4 temporal-steps] of single-trial time-frequency amplitude maps in the high-γ band or those in the α band. A given spectra-temporal feature equals the averaged time-frequency amplitude in a corresponding bin. All SVM analyses were performed in MATLAB using functions from the LIBSVM toolbox (https://www.csie.ntu.edu.tw/~cjlin/libsvm/). For each site, the decoding accuracy of the high-γ band model and that of the α band model were obtained, respectively. To test the significance of the decoding accuracy, we compared the observed decoding accuracy with a null distribution generated by permutation (swapping trial labels before SVM training) 1000 times.

### Statistical analyses

Statistical analyses were performed with IBM SPSS Statistics 22 software (SPSS, Chicago, IL, USA). Paired *t*-tests, independent-samples *t*-tests, chi-square tests, binary logistic regressions, analyses of variance (ANOVAs), and post hoc comparisons (with *Bonferroni* corrections) tests were conducted. The null hypothesis rejection level was set at 0.05.

## Acknowledgments

This work was supported by the National Science and Technology Innovation 2030 Major Program (2022ZD0204802, 2022ZD0204804) and the National Natural Science Foundation of China (31930053, 32171039).

## Author contributions

F.F., and W.Q. conceived the experiments. L.L., X.N, W.J., Y. -R.L., G.-P.C and W.Q. collected the data. R. J. and G.-M.L. performed the surgeries. L.L., Y. -R.L., and W.Q. analyzed the data. F.F., and W.Q. wrote and revised the paper with input from all authors.

