## Supplemental Tables for "Neural response properties predict perceived contents and locations elicited by intracranial electrical stimulation of human auditory cortex": 2. [AudiES] SuppM230506.pdf

**Supplemental Table 1. Patients' demographic and clinical data**

| Patient# | Age | Sex | Implanted hemisphere | Count of selected sites |
|---|---|---|---|---|
| 121 | 24 | F | L | 4 |
| 172 | 30 | F | R | 4 |
| 206 | 26 | M | L | 3 |
| 207 | 28 | M | R | 3 |
| 220 | 26 | M | L | 3 |
| 229 | 28 | M | R | 8 |
| 236 | 18 | M | R | 2 |
| 240 | 29 | F | L | 1 |
| 243 | 27 | M | R | 4 |
| 263 | 28 | M | L | 5 |
| 272 | 31 | M | R | 3 |
| 276 | 20 | M | L | 1 |
| 280 | 23 | M | L, R | 9 |
| 281 | 31 | F | L | 2 |
| 282 | 27 | F | L | 5 |
| 290 | 26 | M | L, R | 4 |
| 361 | 27 | F | R | 3 |
| 365 | 27 | M | L | 3 |
| 370 | 25 | F | L, R | 11 |
| 373 | 15 | M | R | 6 |
| 381 | 27 | F | L | 1 |
| 386 | 29 | M | L, R | 2 |
| 409 | 21 | M | R | 4 |
| 412 | 29 | M | L | 4 |
| 413 | 43 | F | L | 6 |
| 414 | 31 | M | L | 5 |
| 419 | 34 | N | R | 5 |
| 421 | 13 | F | L | 2 |
| **Sum** | | | | 113 |

F, female; M, male; L, left; R, right;

**Supplemental Table 2. Subjective reports of iES-elicited auditory percepts**

| Site ID (Anode) | Brain area | Hemi-sphere | Elicitation threshold (mA) | Subjective reports | Perceived location | Perceived content | |
|---|---|---|---|---|---|---|---|
| P121E03 | HG | L | 2.0 | Sounds in the right ear appear. | contralateral | hallucination | simple sound |
| P121E04 | HG | L | 2.0 | Sounds in the right ear appear. | contralateral | hallucination | simple sound |
| P121E08 | mSTG | L | 3.0 | Sounds in the right ear appear. | contralateral | hallucination | simple sound |
| P121E09 | mSTG | L | 3.0 | Sounds in the right ear appear. | contralateral | hallucination | simple sound |
| P172DD04 | HG | R | 1.0 | A sound appears. | no report | hallucination | simple sound |
| P172DD08 | mSTG | R | 2.0 | A sound appears, louder than before (DD07). | no report | hallucination | simple sound |
| P172DD09 | mSTG | R | 2.0 | A sound appears, lighter than before (DD08). | no report | hallucination | simple sound |
| P172DD10 | mSTG | R | 1.0 | A sound appears, lighter than before (DD09). | no report | hallucination | simple sound |
| P206G04 | HG | L | 3.0 | A "cheeping" sound in the right ear appears. | contralateral | hallucination | simple sound |
| P206G05 | HG | L | 2.0 | A "buzzing" sound in the right ear appears. | contralateral | hallucination | simple sound |

| | | | | | | | |
|---|---|---|---|---|---|---|---|
| P206G07 | HG | L | 3.0 | A "chirp" sound in the right ear appears. | contralateral | hallucination | simple sound |
| P207GG05 | HG | R | 2.0 | A "buzzing" sound in the left ear appears. | contralateral | hallucination | simple sound |
| P207GG06 | HG | R | 2.0 | A "buzzing" sound in the left ear appears. | contralateral | hallucination | simple sound |
| P207GG09 | mSTG | R | 5.0 | A sound in the left ear appears. | contralateral | hallucination | simple sound |
| P220F04 | HG | L | 2.0 | Cannot hear external sound in the right ear. Feels isolated and can only hear his own voice. | contralateral | illusion | suppression |
| P220F05 | HG | L | 3.0 | Cannot hear external sound in the right ear. Feels isolated and can only hear his own voice. | contralateral | illusion | suppression |
| P220F06 | HG | L | 3.0 | Cannot hear external sound in the right ear. Feels isolated and can only hear his own voice. | contralateral | illusion | suppression |
| P229GG03 | HG | R | 3.0 | Cannot hear his own voice. Experiences palpitations. | contralateral | illusion | suppression |
| P229GG04 | HG | R | 1.0 | Cannot hear his own voice. Cannot hear others' voices clearly. Experiences changes in pitch of voices. Experiences palpitations. | no report | illusion | suppression |
| P229GG05 | HG | R | 1.0 | "Cheep" sounds in both ears appear. Experiences palpitations. | bilateral | hallucination | simple sound |

| | | | | | | | |
|---|---|---|---|---|---|---|---|
| P229GG06 | HG | R | 1.0 | "Cheep" sounds in both ears appear. Experiences palpitations. | bilateral | hallucination | simple sound |
| P229GG07 | pSTG | R | 1.0 | Feels like listening through a layer of paper. | no report | illusion | suppression |
| P229GG08 | pSTG | R | 2.0 | "Cheep" sounds in both ears appear. Experiences palpitations. | bilateral | hallucination | simple sound |
| P229GG09 | pSTG | R | 5.0 | Experiences tinnitus in both ears. The sound gradually becomes quiet. | bilateral | hallucination | simple sound |
| P229GG10 | pSTG | R | 3.0 | Experiences tinnitus in both ears. The sound gradually becomes quiet. Experiences palpitations as well. | bilateral | hallucination | simple sound |
| P236GG05 | HG | R | 1.0 | A "dong dong" sound in the left ear appears. External sounds become quieter. | contralateral | illusion | suppression |
| P236GG06 | HG | R | 1.0 | A "dong dong" sound in the left ear appears. External sounds become quieter. | contralateral | illusion | suppression |
| P240F04 | HG | L | 2.5 | External sounds echo in the right. Like surround sounds. | contralateral | illusion | echo |
| P243GG04 | HG | R | 1.0 | A "deng deng" sound in the left ear appears. | contralateral | hallucination | simple sound |
| P243GG05 | HG | R | 1.0 | A "buzzing" sound with a short duration in the left ear appears. | contralateral | hallucination | simple sound |
| P243GG06 | HG | R | 1.0 | A "buzzing" sound with a short duration in the left ear appears. | contralateral | hallucination | simple sound |
| P243GG10 | mSTG | R | 1.0 | A sound within the left ear appears. Discomfort in the throat. | contralateral | hallucination | simple sound |

| | | | | | | | |
|---|---|---|---|---|---|---|---|
| P263G04 | HG | L | 2.0 | A "buzzing" sound in the right ear appears. The sound gradually becomes quiet. | contralateral | hallucination | simple sound |
| P263G05 | HG | L | 2.5 | A continuous "buzzing" sound in the right ear appears. | contralateral | hallucination | simple sound |
| P263G06 | HG | L | 4.0 | A "buzzing" sound moves from the head to the right ear and disappears at the right ear. | contralateral | hallucination | simple sound |
| P263G07 | pSTG | L | 4.0 | A "buzzing" sound in both ears appears. | bilateral | hallucination | simple sound |
| P263G08 | pSTG | L | 4.0 | A "buzzing" sound in both ears appears. | bilateral | hallucination | simple sound |
| P272GG05 | HG | R | 1.5 | A sound like the wind in the left ear appears. | contralateral | hallucination | simple sound |
| P272GG06 | HG | R | 2.0 | A sound like the wind from the left appears. | contralateral | hallucination | simple sound |
| P272GG07 | HG | R | 4.5 | A sound like the wind in the left ear appears. | contralateral | hallucination | simple sound |
| P276G03 | HG | L | 3.0 | A wave-like sound in the right ear appears. | contralateral | hallucination | simple sound |
| P280GG04 | HG | R | 1.5 | External sounds echo in the left. Echoes gradually become quiet. | contralateral | illusion | echo |
| P280GG05 | HG | R | 1.5 | External sounds echo in the left. Echoes gradually become quiet. | contralateral | illusion | echo |
| P280GG06 | HG | R | 1.5 | External sounds echo in the left. Echoes gradually become quiet. | contralateral | illusion | echo |

| | | | | | | | |
|---|---|---|---|---|---|---|---|
| P280GG07 | mSTG | R | 1.5 | External sounds echo in the left. Echoes gradually become quiet. | contralateral | illusion | echo |
| P280GG08 | mSTG | R | 2.5 | External sounds echo in the left. Echoes gradually become quiet. | contralateral | illusion | echo |
| P280GG09 | mSTG | R | 4.0 | External sounds echo in the left. Echoes gradually become quiet. | contralateral | illusion | echo |
| P280R04 | HG | L | 1.5 | An "en en" sound in both ears appears from behind. | bilateral | illusion | echo |
| P280R05 | HG | L | 1.5 | A "buzzing" sound with a short duration in both ears appears from behind. | bilateral | hallucination | simple sound |
| P280R06 | HG | L | 1.5 | A "buzzing" sound with a short duration in both ears appears from behind. | bilateral | hallucination | simple sound |
| P281G05 | HG | L | 3.0 | A "buzzing" sound from the right side appears. Feels discomfort. | contralateral | hallucination | simple sound |
| P281G06 | HG | L | 3.0 | Sound echoes in the right ear. Feels discomfort. | contralateral | illusion | echo |
| P282G04 | HG | L | 2.0 | A continuous "ah" sound in the right ear appears. | contralateral | hallucination | simple sound |
| P282G05 | HG | L | 1.5 | Her own voice echoes. | no report | illusion | echo |
| P282G06 | HG | L | 1.5 | Her own voice echoes. | no report | illusion | echo |
| P282G07 | HG | L | 1.5 | Her own voice echoes. | no report | illusion | echo |
| P282G08 | HG | L | 1.5 | Her own voice echoes. | no report | illusion | echo |
| P290G05 | HG | L | 2.0 | Cannot hear anything in both ears. | bilateral | illusion | suppression |
| P290G06 | HG | L | 1.5 | Cannot hear anything in both ears. | bilateral | illusion | suppression |
| P290G07 | mSTG | L | 1.5 | Cannot hear anything in both ears. | bilateral | illusion | suppression |
| P290G08 | mSTG | L | 1.5 | Cannot hear anything in both ears. | bilateral | illusion | suppression |

| | | | | | | | |
|---|---|---|---|---|---|---|---|
| P361N06 | HG | R | 1.0 | A sound like the electric current in the left ear appears. Unable to describe details. | contralateral | hallucination | simple sound |
| P361N07 | HG | R | 1.0 | A sound like the electric current in the left ear appears. Unable to describe details. | contralateral | hallucination | simple sound |
| P361N08 | HG | R | 1.0 | A sound like the electric current in the left ear appears. Unable to describe details. | contralateral | hallucination | simple sound |
| P365S03 | HG | L | 1.5 | A "buzzing" sound in the right ear appears. The sound then moves to the left ear. | bilateral | hallucination | simple sound |
| P365S04 | HG | L | 1.5 | A sound appears. The sound then moves to the posterior side. | no report | hallucination | simple sound |
| P365S05 | HG | L | 1.5 | A sound like a cicada chirping in both ears appears. | bilateral | hallucination | simple sound |
| P370G04 | HG | L | 1.5 | A shrilling sound in the right ear appears. | contralateral | hallucination | simple sound |
| P370G05 | HG | L | 1.0 | A continuous "ta ta" sound in the right ear appears. | contralateral | hallucination | simple sound |
| P370G06 | HG | L | 1.0 | A continuous "ta ta" sound in the right ear appears. | contralateral | hallucination | simple sound |
| P370G07 | HG | L | 1.5 | A continuous "ta ta" sound from the right-middle side appears. | contralateral | hallucination | simple sound |
| P370G08 | HG | L | 1.5 | A continuous "ta ta" sound from the right-middle side appears. | contralateral | hallucination | simple sound |
| P370G09 | pSTG | L | 6.0 | A sound like the wind from the right side appears. | contralateral | hallucination | simple sound |

| | | | | | | | |
|---|---|---|---|---|---|---|---|
| P370GG05 | HG | R | 2.0 | A "sizzling" sound in the left ear appears. | contralateral | hallucination | simple sound |
| P370GG06 | HG | R | 2.0 | A "sizzling" sound in the left ear appears. | contralateral | hallucination | simple sound |
| P370GG07 | HG | R | 3.0 | A sound with pitch perception appears. | no report | hallucination | simple sound |
| P370GG08 | pSTG | R | 4.0 | A sound with rhythm appears. | no report | hallucination | simple sound |
| P370GG09 | pSTG | R | 5.0 | A sound with pitch perception appears. | no report | hallucination | simple sound |
| P373GG04 | HG | R | 1.0 | A "biu biu" sound like a water pistol appears in the left ear. The sound gradually becomes quiet. | contralateral | hallucination | simple sound |
| P373GG05 | HG | R | 1.0 | A "biu biu" sound like a water pistol appears in the left ear. The sound gradually becomes quiet. | contralateral | hallucination | simple sound |
| P373GG06 | HG | R | 1.5 | A "biu biu" sound like a water pistol appears in the left ear. The sound gradually becomes quiet. | contralateral | hallucination | simple sound |
| P373GG07 | HG | R | 2.0 | A "biu biu" sound like a water pistol appears in the left ear. The sound gradually becomes quiet. | contralateral | hallucination | simple sound |
| P373GG08 | HG | R | 3.5 | A "biu biu" sound like a water pistol appears in the left ear. The sound gradually becomes quiet. | contralateral | hallucination | simple sound |

| | | | | | | | |
|---|---|---|---|---|---|---|---|
| P373GG09 | pSTG | R | 5.5 | A "biu biu" sound like a water pistol appears in the left ear. The sound gradually becomes quiet. | contralateral | hallucination | simple sound |
| P381G05 | HG | L | 2.5 | A "buzzing" sound in the right ears appears. | contralateral | hallucination | simple sound |
| P386FF03 | mSTG | R | 2.0 | Hears someone singing in the left ear. Unable to describe details. | contralateral | hallucination | song |
| P386FF04 | mSTG | R | 2.0 | Hears someone singing a song in his head. Unable to describe details. | no report | hallucination | song |
| P409VV01 | HG | R | 1.0 | A "rustling" sound in the left ear appears. | contralateral | hallucination | simple sound |
| P409VV02 | HG | R | 1.0 | A "rustling" sound in the left ear appears. | contralateral | hallucination | simple sound |
| P409VV03 | HG | R | 1.0 | A "rustling" sound in the left ear appears. The sound is lighter. | contralateral | hallucination | simple sound |
| P409VV04 | pSTG | R | 1.0 | A "rustling" sound in the left ear appears. The sound is lighter. | contralateral | hallucination | simple sound |
| P412G03 | HG | L | 3.5 | The speech sound in the right ear echoes. | contralateral | illusion | echo |
| P412G04 | HG | L | 1.0 | A sound like car-roaring in the right ear appears. | contralateral | hallucination | simple sound |
| P412G05 | HG | L | 1.0 | The speech sound in the right ear echoes. | contralateral | illusion | echo |
| P412G07 | pSTG | L | 2.0 | The speech sound in the right ear echoes. | contralateral | illusion | echo |
| P413G04 | HG | L | 2.0 | An "emmm emmm" sound in the right ear appears. | contralateral | hallucination | simple sound |
| P413G05 | HG | L | 2.5 | Hears a familiar melody in the right ear. | contralateral | hallucination | samiliar music |

| | | | | | | | |
|---|---|---|---|---|---|---|---|
| P413G06 | HG | L | 2.5 | An "emmm emmm" sound inside the head appears. | bilateral | hallucination | simple sound |
| P413G08 | pSTG | L | 2.5 | Sound echoes in the head. | bilateral | illusion | echo |
| P413G09 | pSTG | L | 3.0 | Sound echoes in the head. | bilateral | illusion | echo |
| P413G10 | pSTG | L | 3.0 | Sound echoes in the head. | bilateral | illusion | echo |
| P414G05 | HG | L | 2.5 | Sound echoes in the right ear. Voice of his own becomes louder. | contralateral | illusion | echo |
| P414G06 | HG | L | 2.0 | Sound echoes in the right ear. Voice of his own becomes louder. | contralateral | illusion | echo |
| P414G07 | pSTG | L | 1.5 | Sound echoes in the right ear. Voice of his own becomes louder. | contralateral | illusion | echo |
| P414G08 | pSTG | L | 2.5 | Sound echoes in the right ear. Voice of his own becomes louder. | contralateral | illusion | echo |
| P414G09 | pSTG | L | 4.0 | Sound echoes in the right ear. Voice of his own becomes louder. | contralateral | illusion | echo |
| P419GG04 | HG | R | 1.5 | A cicada chirping in both ears appears. Sound in the left ear is louder than in the right ear | bilateral | hallucination | simple sound |
| P419GG05 | HG | R | 1.5 | A cicada chirping in the left ear appears. | contralateral | hallucination | simple sound |
| P419GG06 | HG | R | 2.0 | A cicada chirping in the left ear appears. | contralateral | hallucination | simple sound |
| P419GG07 | mSTG | R | 2.5 | A cicada chirping appears. | no report | hallucination | simple sound |
| P419GG08 | HG | R | 1.0 | A "buzzing" sound appears. | no report | hallucination | simple sound |

| | | | | | | | |
|---|---|---|---|---|---|---|---|
| P421G02 | HG | L | 1.0 | A noise-like sound in the right ear appears. Unable to describe further details. | contralateral | hallucination | simple sound |
| P421G03 | mSTG | L | 1.0 | A cicada chirping in the right ear appears. | contralateral | hallucination | simple sound |

*L = left; R = right; HG = Heshl's gyrus; mSTG = middle superior temporal gyrus; pSTG = posterior superior temporal gyrus; iES = intracranial electrical stimulation. iES parameters: duration = 5 s, frequency = 50 Hz, pulse width =10 μS.
